# DRAM1 promotes antibacterial autophagy and lysosomal delivery of Mycobacterium marinum in macrophages

**DOI:** 10.1101/2022.10.25.513660

**Authors:** Adrianna Banducci-Karp, Jiajun Xie, Sem A. G. Engels, Christos Sarantaris, Monica Varela, Annemarie H. Meijer, Michiel van der Vaart

## Abstract

Damage-Regulated Autophagy Modulator 1 (DRAM1) is an infection-inducible membrane protein, whose function in the immune response is incompletely understood. Based on previous results in a zebrafish infection model, we have proposed that DRAM1 is a host resistance factor against intracellular mycobacterial infection. To gain insight into the cellular processes underlying DRAM1-mediated host defence, here we studied the interaction of DRAM1 with Mycobacterium marinum in murine RAW264.7 macrophages. We found that shortly after phagocytosis, DRAM1 localised in a punctate pattern to mycobacteria, which gradually progressed to full DRAM1 envelopment of the bacteria. Within the same time frame, DRAM1-positive mycobacteria colocalised with the LC3 marker for autophagosomes and LysoTracker and LAMP1 markers for (endo)lysosomes. Knockdown analysis revealed that DRAM1 is required for recruitment of LC3 and for acidification of mycobacteria-containing vesicles. A reduction in the presence of LAMP1 further suggested reduced fusion of lysosomes with mycobacteria-containing vesicles. Finally, we show that DRAM1 knockdown impairs the ability of macrophages to control mycobacterial infection. Together, these results support that DRAM1 promotes the trafficking of mycobacteria through the degradative (auto)phagolysosomal pathway. Considering its prominent effect on host resistance to intracellular infection, DRAM1 is a promising target for therapeutic modulation of the microbicidal capacity of macrophages.

## 1. Introduction

Innate immune cells, including macrophages, are the gatekeepers of the immune system: they ingest microbes into internal compartments called phagosomes, they mount a range of cell-autonomous defence responses, and they alert other components of the immune system. A key cell-autonomous defence response is the phagolysosomal pathway, which delivers microbes through a phagosome maturation process to lysosomes. Upon lysosomal fusion, microbes are exposed to an acidic environment, antimicrobial peptides, and degradative enzymes [1]. However, several intracellular pathogens are able to withstand the phagocyte defences because they are able to inhibit the maturation of phagosomes, adapt to the hostile environment of lysosomes, or breach the integrity of the phagosome membrane and invade the cytosol [2]. Invasion of the cytosol exposes these pathogens to a secondary innate immune defence process, termed autophagy, which serves as a mechanism to preserve cellular homeostasis [3]. In the anti-microbial autophagy response, also known as xenophagy, selective autophagy receptors recognize microbes marked for proteasomal degradation by ubiquitin coating [4]. These receptors then recruit the autophagy machinery through interaction with microtubule-associated proteins 1 light chain 3 (hereafter LC3). As a result, microbes are captured inside double membrane vesicles, called autophagosomes, which subsequently fuse with lysosomes to degrade its contents. Similar as for the phagolysosomal route, several pathogens have evolved mechanisms to inhibit the autolysosomal route at various points in the process [5].

The genus Mycobacterium comprises several human pathogens that are able to subvert both the phagolysosomal and autolysosomal pathways to facilitate their intracellular survival in macrophages [6–9]. The most notorious mycobacterial species is Mycobacterium tuberculosis (Mtb), which causes 10 million cases of tuberculosis (TB) and close to 1.5 million deaths every year [10]. However, the medical importance of mycobacteria extends beyond Mtb to a large collection of nontuberculous mycobacteria (NTM). For example, Mycobacterium avium lung infections are associated with poor clinical outcome and are notably on the rise [11,12]. Another example of NTMs is Mycobacterium marinum (Mm), known for outbreaks of fish TB-like infection in aquaculture as well as for skin infections in humans [13]. Mm is closely related to Mtb, sharing 3000 orthologous proteins with an average amino acid identity of 85% [14]. This similarity includes the RD1 virulence locus encoding for the type VII secretion system ESX-1 and its secreted proteins, ESAT-6 and CFP-10 [15]. In both Mm and Mtb, the bacteria require the ESX-1 system to evade phagolysosomal degradation and invade the cytosol [16]). As a countermeasure, ESX-1 competent Mm or Mtb are targeted by xenophagy [17–19]; however, it remains unknown to what extent xenophagy provides host protection during clinical infection as several studies report that mycobacteria possess virulence mechanisms to inhibit autophagy-mediated degradation [20].

DRAM1, for Damage-regulated autophagy modulator 1, is the founding member of a family of five stress-inducible autophagy regulators [21,22]. DRAM1 functions as a target of p53-signalling in the UV-damage response and as a target of NF-κB-signalling during Mm and Mtb infection [18,21]. In addition, DRAM2 has been implicated in defence against mycobacteria, whereas DRAM3, 4, and 5 have not been linked to infectious diseases [23–26]. We have shown previously that DRAM1 provides a link between pathogen recognition and activation of autophagy, as its induction during mycobacterial infection was found to be dependent on MyD88 and NFκB, central mediators of the Toll-like receptor pathway [18]. Using a zebrafish model for Mm infection, we showed that knockdown or mutation of the zebrafish ortholog of DRAM1 causes hypersusceptibility to infection, while its overexpression increases host resistance against Mm [18,27]. We also obtained evidence that the zebrafish DRAM1 ortholog promotes the association of Mm with LC3 and increases lysosomal acidification of Mm-containing compartments [18,27]. However, due to the lack of antibody tools for zebrafish, we have not been able to study how DRAM1 colocalises with mycobacteria, autophagosomes and lysosomes during the course of infection.

In order to study DRAM1 localization and function during mycobacterial infection in a relevant mammalian cell type, we used the mouse RAW264.7 macrophage cell line, which has been frequently employed for Mtb autophagy studies [28–30]. Mm infection of this cell line leads to rapid permeabilisation of phagosomes and subsequent ubiquitination of bacteria, marking them as targets for xenophagy [31]. Here, we show the progression of the colocalisation of DRAM1 with Mm over the course of infection and study this in relation to LC3, LysoTracker and LAMP1 patterns as markers for autophagosomes and (endo)lysosomes. In addition, we generated DRAM1 knockdown macrophage cell lines to show that DRAM1 knockdown reduces both the trafficking of LC3 to Mm and the acidification of Mm-containing vesicles. Furthermore, upon DRAM1 knockdown, the Mm infection rate increased. These results support the central role of DRAM1 in host resistance to intracellular infection by promoting the trafficking of mycobacteria in the (auto)phagolysosomal pathway.

## 2. Materials and Methods

### 2.1 Macrophage cell culture

The RAW 264.7 macrophage cell line was maintained in Dulbecco’s Modified Eagle’s Medium (DMEM) high glucose (Sigma-Aldrich, D6546) with 10% fetal calf serum (FCS, F2442) (Gibco) at 37°C in a 5% CO2 atmosphere. The day before Mm infection, RAW 264.7 cells were seeded on sterile coverslips inserted in 12 well plates (2.5×105 cells/well for the 0-150 mpi Mm infection experiments or 4×105 cells/well for the Mm infection rate experiment) and incubated overnight at 37°C with 5% CO2.

### 2.2 Mm culture and infection experiments

The day before cell infection, Mm M-strain, fluorescently labelled with mWasabi [32], was cultured in Difco Middlebrook 7H9 medium (Becton Dickinson, BD271310) with 10% BBL™ Middlebrook albumin-dextrose-catalase (Becton Dickinson, 211887) and 50 μg/ml hygromycin (Sigma-Aldrich, SC-506168A) at 28.5°C with in a static incubator. On the day of infection, Mm was washed twice with PBS and optical density (OD) was measured at 600nm. OD600nm of 1 is equal to 108 Mm/mL [33]. RAW 264.7 cells were then infected with a multiplicity of infection (MOI) of 10, immediately centrifuged at 500 g for 10 minutes to synchronise infection, as described in [34], and incubated at 32°C with 5% CO2. After 30 minutes, cells were washed twice with PBS to remove extracellular Mm and then incubated with LysoTracker-enriched medium for up to 150 min at 32°C to obtain samples for immunofluorescence at 30 min intervals. At each 30-min interval, cells were fixed with Pierce™ 4% formaldehyde (PFA) (Thermo Scientific, 28908) for 20 minutes at room temperature (RT). To analyse infection rate, cells were infected at a MOI of 10 without centrifugation and incubated at 32°C. After 1 hour, cells were either fixed with PFA or washed with PBS twice to remove extracellular bacteria and then incubated with fresh medium for 6 hours.

### 2.3 Immunofluorescence

After PFA fixation, cells were washed with PBS and permeabilised with 0.2% Saponin (Sigma-Aldrich, 47036) in PBS for 15 min, before blocking with 1% bovine serum albumin (BSA) (Sigma-Aldrich, A4503) in PBS for 1 hour at RT. Next, cells were incubated with the appropriate primary antibody, specifically DRAM1 (1:500) (Invitrogen, OSD00007G), LC3 (1:500) (Novus Biologicals, NB100-2331), or LAMP1 (Abcam, ab24170) for 1 hour at RT. For single immunostaining, goat-anti-rabbit Alexa Fluor™ Plus 647 (1:1000) (Invitrogen, A-21245) was added to detect the primary antibodies. Between each step, cells were washed four times with PBS for 5 min each. For double immunostaining, after permeabilisation and blocking with 1% BSA, cells were incubated with DRAM1 primary antibody (1:250) for 2 hours and then washed with PBS 3×10min. Then, the goat-anti-rabbit Fab 647 (1:50) (Sanbio, 111-607-003) was incubated overnight. The next day, cells were washed 6×10 min. Cells were incubated with Fab secondary antibody again for 2 hours at RT. After washing, cells were incubated with the LC3 primary antibody (1:250) for 1.5 hours and afterwards washed for 3×10 min. Lastly, cells were incubated with the second secondary antibody goat-anti-rabbit Alexa Fluor™ 568 (1:1000) (Invitrogen, A-11011) for 2 hours at RT and then washed for 4×5min. Cover slips were mounted with Prolong Gold Antifade Reagent (Invitrogen, P36966) onto glass slides for imaging.

### 2.4 LysoTracker labelling

Acidic vesicles were visualised by incubating RAW 264.7 cells in medium enriched with LysoTracker™ Red DND-99 (1:2500) (Invitrogen, L7528) for at least 1 hour prior to fixation at 37°C [35]. For non-infected cells, LysoTracker staining was performed for 30 min before fixation.

### 2.5 DRAM1 shRNA knockdown

To knockdown DRAM1, three short hairpin RNAs (shRNAs) against DRAM1 (NM_027878) from the Mission library (Sigma-Aldrich) were used. pLKO.1-puro shRNA constructs were transduced into RAW 264.7 cells. The day before transduction, 1×106 RAW 264.7 cells were seeded per T25 flask in DMEM-high glucose with 10% FCS and incubated overnight at 37°C with 5% CO2. 24 hours later, cells were transduced at a MOI of 4 or 8, supplemented with 8 μg/ml polybrene (Sigma-Aldrich, TR1003), and incubated at 37°C with 5% CO2. Medium was refreshed 24 hours after transduction. After 48 hours of transduction, 3 μg/ml puromycin (Gibco, A1113803) was added. Every 2-3 days, the medium was replaced until cells were 80-90% confluent. To isolate single clones with successful DRAM1 knockdown, cells from each condition were diluted in DMEM-high glucose with 10% FCS and 3 μg/ml puromycin to achieve 1 cell/well in a 96 well plate. The following day, wells containing 1 cell were marked. Once marked wells reached 80-90% confluency, cells were transferred to 12-well plates and subsequently to T25 flasks. Antibiotic pressure was removed from the medium to confirm stability of DRAM1 shRNA integration into cells.

### 2.6 Quantitative PCR

Cell pellets containing 1×106 cells were dissolved in TRIzol (Invitrogen, 15596018) and then chloroform (Supelco^®^, 1.07024.2500). After centrifuging at 12,000g for 15 min at 4°C, the aqueous phase containing the RNA was transferred to a new tube. One volume of 70% ethanol was added and then transferred to a column from the RNeasy mini kit (Qiagen, 74104). RNA isolation was performed according to the manufacturer’s protocol, which included on-column DNAse digestion. 500 ng of each RNA sample, measured with the Nanodrop (Thermofisher), was converted to cDNA with the C1000 Touch Thermal Cycler using the iScript cDNA Synthesis Kit according to the manufacturer’s protocol (Bio-Rad, 1708890), with the following modifications: 1 μl iScript Reverse Transcriptase was modified to 0.5 μl. Samples were then subjected to quantitative PCR (qPCR) using the CFX96 Real-Time System (Bio-Rad) with the iTaq™ Universal SYBR^®^ Green Supermix (Bio-Rad, 1725271). The program was set up as follows: initial denaturation at 95°C for 3 minutes then 40 cycles of 95°C for 15 seconds (denaturation) and 60°C for 30 seconds (annealing and extension). A melting curve was included to test for PCR product purity: 55-95°C in 0.5°C increments. DRAM1 primers (Sigma-Aldrich) used are as follows: Forward: 5’-CCAGCTTCTTGGTCCGACG-3’, Reverse: ‘5-GGGAGAAAGGGGTTGACGTG-3’. GAPDH was used as the housekeeping gene: Forward: 5’-ATGGTGAAGGTCGGTGTGAA-3’, Reverse: ‘5-CTGGAACATGTAGACCATGT-3’.

### 2.7 Western blot

Cells were harvested, lysed in RIPA buffer (Cell Signalling, 9806) containing a protein inhibitor cocktail (Roche, 11873580001), and centrifuged at 4°C for 10 min at 12,000 g/min. Western blotting was performed using 15% polyacrylamide gel for LC3 and 10% for polyacrylamide gels for DRAM1 and LAMP1, followed by protein transfer to commercial PVDF membranes (Bio-Rad, 1704156). Membranes were blocked with 3% BSA in 1x Tris buffered saline (TBS) solution with 0.1% Tween-20 (TBST) before incubating with primary and secondary antibodies: polyclonal rabbit anti DRAM1 (1:1000) (Aviva systems biology, ARP47432-P050), GAPDH (1:1000)(D16H11), LC3 (1:1000) (Novus Biologicals, NB100-2331), LAMP1 (1:1000) (Abcam, ab24170), Antirabbit IgG, HRP-Linked Antibody (1:1000) (Cell Signaling, 7074S). Digital images were acquired using the Bio-Rad Universal Hood II imaging system (720BR/01565 UAS). Band intensities were quantified with ImageJ and values were normalized to GAPDH as a loading control.

### 2.8 Confocal laser scanning microscopy

RAW 264.7 cells stained by immunofluorescence and/or LysoTracker dye were imaged with a TCS SP8 confocal microscope (Leica) using a 63x oil immersion objective (NA: 1.40). Images were processed using the ImageJ software.

### 2.9 Statistical analyses

Analysis of DRAM1-Mm colocalisation involved the characterisation into four categories (none, membrane, punctate, luminal) based on visual inspection of the images. ‘None’: no colocalisation between DRAM1 and Mm. ‘Membrane’: DRAM1 staining surrounds Mm in a ring. ‘Punctate’: DRAM1 colocalises with Mm but not along Mm’s entire length. ‘Luminal’: DRAM1 completely colocalises with Mm inside the lumen of a Mm-containing vesicle. The same definitions were used to quantify LC3-Mm and LAMP1-Mm colocalisations, where we note that the membrane pattern was not observed for LC3-Mm. Mm clusters fully colocalizing with LysoTracker were also quantified. Fluorescent intensity and particles were analysed in ImageJ. Logistic regression and pairwise comparison with Turkey correction performed in R was used to analyse statistical differences between knockdown and control groups (*p<0.05; **p<0.01; ***p<0.001; ****p<0.0001). All graphs were made in GraphPad Prism8 (mean ± SD).

## 3. Results

### 3.1 DRAM1 colocalises with acidified Mm-containing vesicles

To study the localization of DRAM1 in response to Mm infection in RAW264.7 macrophages, we performed immunofluorescence staining. We first determined the localisation pattern of DRAM1 in non-infected conditions. We observed that DRAM1 concentrates at certain areas in the vicinity of the plasma membrane. In these highly concentrated regions, DRAM1-positive circular structures of various sizes were occasionally observed (Fig.1A). In addition, we observed that DRAM1 localised to subcellular vesicles. Based on co-staining with LysoTracker, these vesicles were often acidic (Fig.1B), in agreement with the previous identification of DRAM1 as a largely lysosomal protein [21,36]. We then continued to study DRAM1 localisation during Mm infection and followed the acidification of Mm-containing vesicles by LysoTracker staining (Fig.2A,B). We determined DRAM1-Mm colocalisation patterns over a time course of 150 min with 30 min intervals. Approximately 40-80% of Mm colocalised with DRAM1 during this time course, with a peak at 120 min (Fig.2C). While DRAM1-Mm colocalised from the start of infection, LysoTracker-Mm colocalisation sharply increased from low levels to approximately 50% at 120 hours (Fig.2B,C). Over the time course, we could distinguish three main categories of DRAM1 signal, which we refer to as membrane, punctate and luminal (Fig.2A). The DRAM1 signals classified in the membrane category did not colocalise with Mm but surrounded Mm-containing vesicles (Fig.2A). This category was the smallest, representing less than 5% of all DRAM1-Mm associations at all time points (Fig.2D). Punctate signals showed as small patches of DRAM1 staining that were closely associated with Mm. These punctate patterns constituted 40-60% of all DRAM1-Mm colocalisation with a relatively stable frequency over the time course (Fig.2D). In the luminal category, DRAM1 staining overlapped fully (Fig.2A, luminal 1) with Mm or contained a weak Mm signal, which might be interpreted as residual staining from degraded bacteria (Fig.2A, luminal 2). These luminal patterns were not detected before 90 min and were most frequent at 120 min, where approximately 30% of DRAM1-Mm associations showed this pattern (Fig.2D). We found that the luminal DRAM1 signals were always LysoTracker-positive (Fig. 2A,E). Mm surrounded by DRAM1 staining (membrane category) was always LysoTracker-negative. LysoTracker staining of punctate DRAM1-Mm patterns increased over time, similar to DRAM1-negative Mm (Fig.2E). In conclusion, increasing overlap between DRAM1 signal and bacteria parallels the acidification of Mm-containing vesicles.

**Figure 1.**
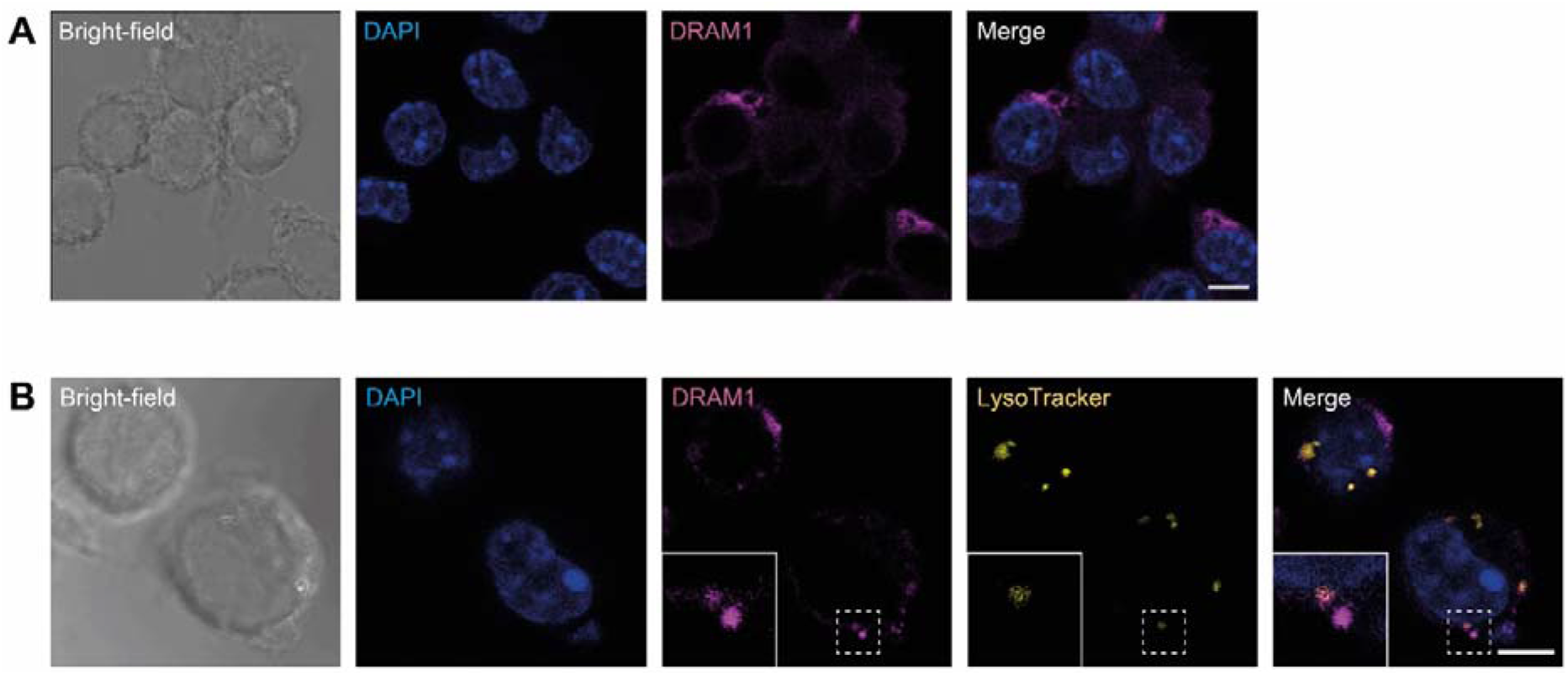
DRAM1 is predominantly localised at the plasma membrane and in acidic vesicles of RAW 267.4 macrophages. (A) DRAM1 immunofluorescence (magenta) with the nucleus stained by DAPI (blue). Scale bars: 5 μm (B) DRAM1 (magenta) colocalisation with acidic vesicles stained by LysoTracker (yellow). The nucleus is stained by DAPI (blue). A region showing a LysoTracker-positive and a LysoTracker-negative DRAM1-stained vesicle is outlined with a dotted box in the merged image and enlarged in the inset. Scale bars: 5 μm.

**Figure 2.**
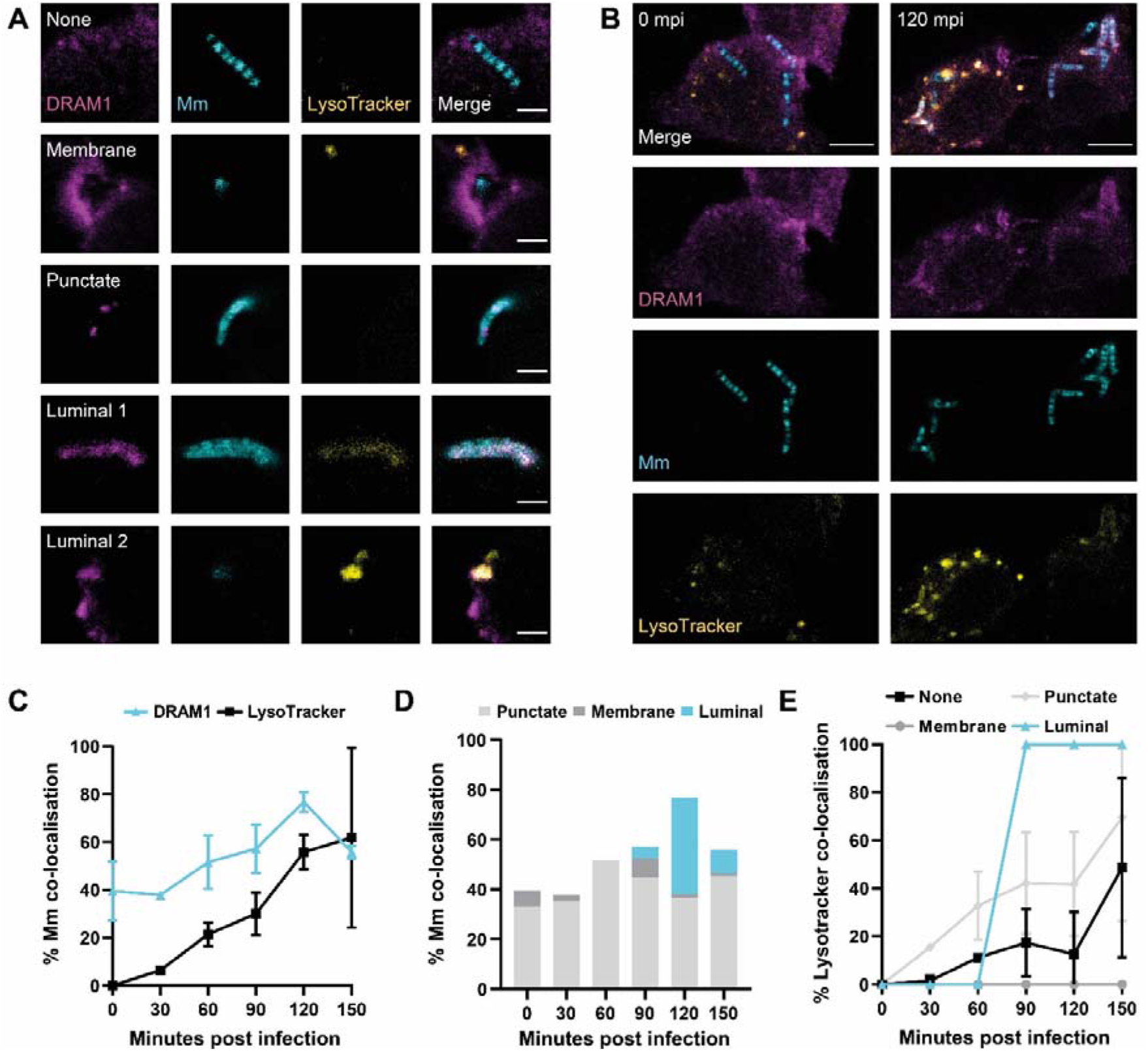
DRAM1 colocalises with acidified mycobacteria-containing vesicles. (A) Representative examples of DRAM1 (magenta) colocalisation patterns with Mm (cyan), referred to as membrane, punctate, and luminal patterns. An example of DRAM1-negative Mm (none) is also shown. Acidification of Mm-containing vesicles is assessed by LysoTracker staining (yellow). Scale bars: 2 μm (B) Representative example of DRAM1 (magenta) colocalisation with LysoTracker (yellow) and Mm (cyan) at 120 min post infection, when luminal colocalisation patterns are most frequent. Scale bars: 5 μm (C) Frequency of Mm colocalisation with DRAM1 or LysoTracker over time. (D) Frequency of Mm colocalisation with DRAM1 in punctate, membrane or luminal patterns over time. (E) Frequency of LysoTracker colocalisation with DRAM1-negative Mm and DRAM1-positive membrane, punctate and luminal Mm patterns over time. Data are accumulated from two independent experiments (n0=71, n30=73, n60=105, n90=98, n120=102, n150=111 bacterial clusters).

### 3.2 Acidified Mm-containing vesicles are positive for the autophagy marker LC3

Having demonstrated the large overlap between DRAM1 and LysoTracker staining, we wished to further assess the identity of the acidified Mm-containing vesicles. Considering that Mm can permeabilise phagosomes to invade the cytosol, Mm may become a substrate for autophagy. Therefore, we investigated the colocalisation between Mm and LC3, a marker for autophagosomes, in combination with LysoTracker staining or DRAM1 staining (Fig.3A,B,C,D). We classified the LC3-Mm associations in 2 categories, punctate and luminal (Fig.3A,E), which were similar to those observed before in DRAM1 immunostaining (Fig.2A). In agreement, for both the punctate and the luminal LC3-Mm patterns, we observed colocalization with DRAM1 (Fig.3C). The luminal LC3-Mm patterns increased in frequency between 0-60 min, after which the punctate and luminal patterns occurred at a relatively stable frequency between 60-150 min (Fig.3E). The luminal signals overlapped with either a strong Mm signal (Fig.3A, luminal 1) or a weaker dispersed Mm signal (Fig.3A, luminal 2), suggesting that bacterial degradation may have occurred in the latter case similar as observed before in DRAM1 immunostaining (Fig.2A, luminal 2). Quantification showed a near complete overlap between luminal LC3 signals and LysoTracker staining, whereas only a subset of punctate LC3 signals were LysoTracker-positive (Fig. 3A,B,D,F). In conclusion, LC3 colocalises with acidified Mm-containing vesicles in a similar pattern to DRAM1.

**Figure 3.**
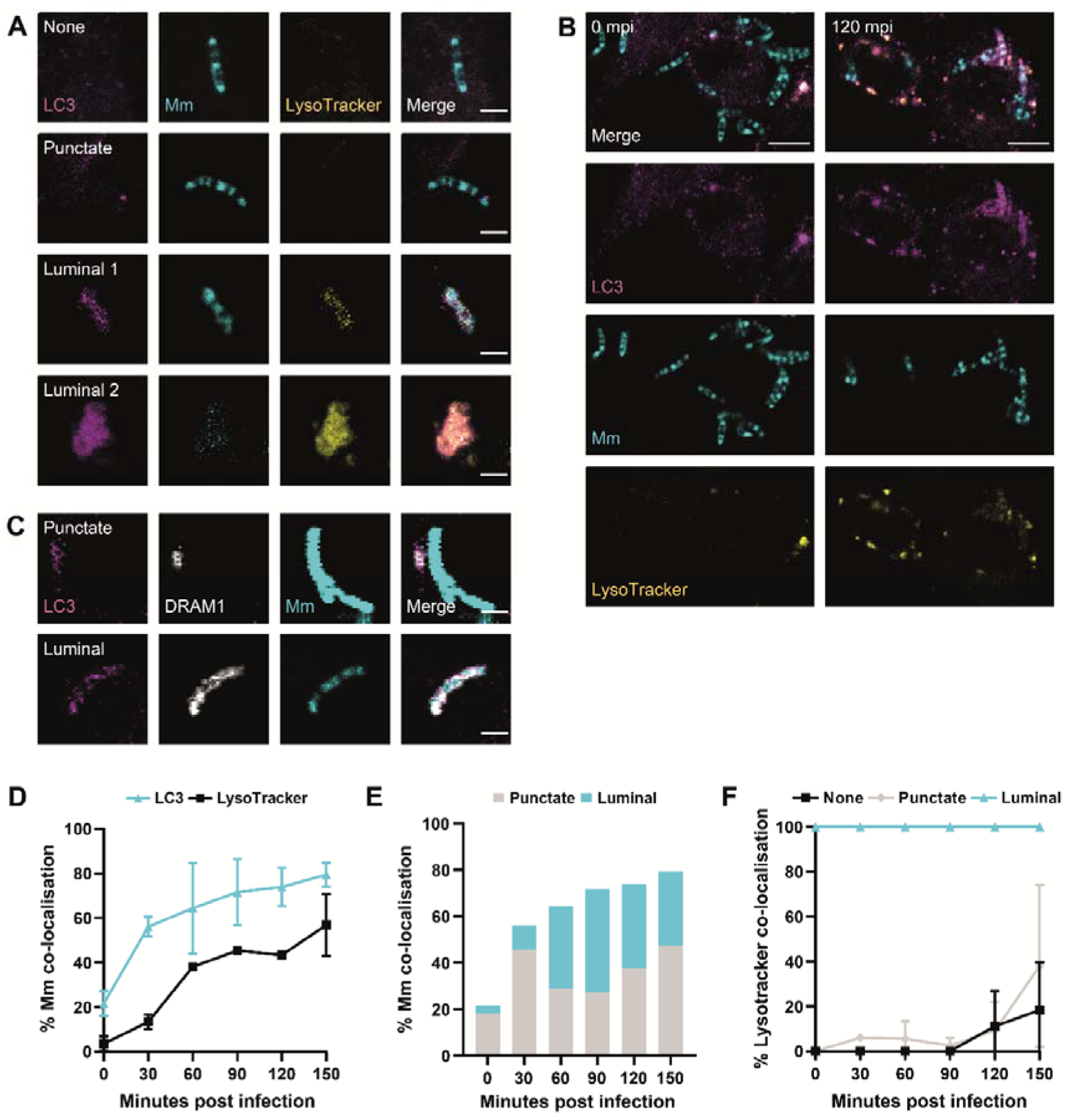
Acidified mycobacteria-containing vesicles are labelled by LC3. (A) Representative examples of LC3 (magenta) colocalisation patterns with Mm (cyan), referred to as punctate and luminal patterns. An example of LC3-negative Mm (none) is also shown. Acidification of Mm-containing vesicles is assessed by LysoTracker staining (yellow). Scale bars: 2 μm (B) Representative example of LC3 (magenta) colocalisation with LysoTracker (yellow) and Mm (cyan) at 120 min post infection, when luminal colocalisation patterns are most frequent. Scale bars: 5 μm (C) Representative examples of double colocalisation of LC3 (magenta) and DRAM1 (grey) with Mm (cyan),. Scale bars: 2 μm (D) Frequency of Mm colocalisation with LC3 or LysoTracker over time. (E) Frequency of Mm colocalisation with LC3 in punctate or luminal patterns over time. (F) Frequency of LysoTracker colocalisation with LC3-negative Mm and LC3-positive punctate and luminal Mm patterns over time. Data are accumulated from two independent experiments (n0=99, n30=129, n60=96, n90=124, n120=133, n150=137 bacterial clusters).

### 3.3 DRAM1 promotes acidification of Mm-containing vesicles

To study the function of DRAM1 during Mm infection in macrophages, we generated three independent DRAM1 shRNA knockdown cell lines by lentiviral transduction and showed that Dram1 mRNA expression levels (Sup Fig.1A), as well as protein levels (Sup Fig.1B), were reduced in each of the three cell lines compared to the control cells. Next, we analysed the effect of DRAM1 knockdown to Mm infection at 120 min post infection, the time point at which DRAM1-Mm colocalisation events were most frequent (Fig.2C). Cells were stained for DRAM1 to further validate the knockdown effect and for LysoTracker to determine the effect of DRAM1 on acidification of Mm-containing vesicles (Fig.4A). As expected, the reduced DRAM1 levels due to shRNA knockdown resulted in less DRAM1-Mm colocalisation events when compared to the control cells (Fig.4B). Furthermore, LysoTracker-Mm colocalisation was reduced, consistent with a role for DRAM1 in promoting vesicle acidification (Fig.4B). While we still observed the same patterns of colocalisation between DRAM1 and Mm in the knockdown cells as in the control, the frequency of luminal patterns was reduced in the knockdown lines (Fig.4C). The colocalisation of these DRAM1-Mm patterns with LysoTracker was also affected; specifically, there was a decrease in LysoTracker staining of the punctate DRAM1-Mm patterns upon DRAM1 knockdown (Fig.4D). In addition, DRAM1-negative Mm clusters also showed a decrease in LysoTracker staining in response to DRAM1 knockdown. In contrast, the luminal DRAM1-Mm patterns always remained LysoTracker positive in the knockdown cell lines. To verify that the decrease in LysoTracker staining of Mm-containing vesicles was not caused by an overall reduction in LysoTracker-positive vesicles due to the knockdown of DRAM1, we performed LysoTracker staining of non-infected cells, which revealed that there were no differences in LysoTracker staining, quantity, or intensity upon DRAM1 knockdown (Supplementary Fig2.A,B,C). Taken together, DRAM1 knockdown reduced acidification of Mm-containing vesicles with DRAM1-negative or punctate DRAM1 patterns. However, when the remaining DRAM1 protein levels in the knockdown lines led to luminal colocalisation with Mm, these events were always associated with acidification.

**Figure 4.**
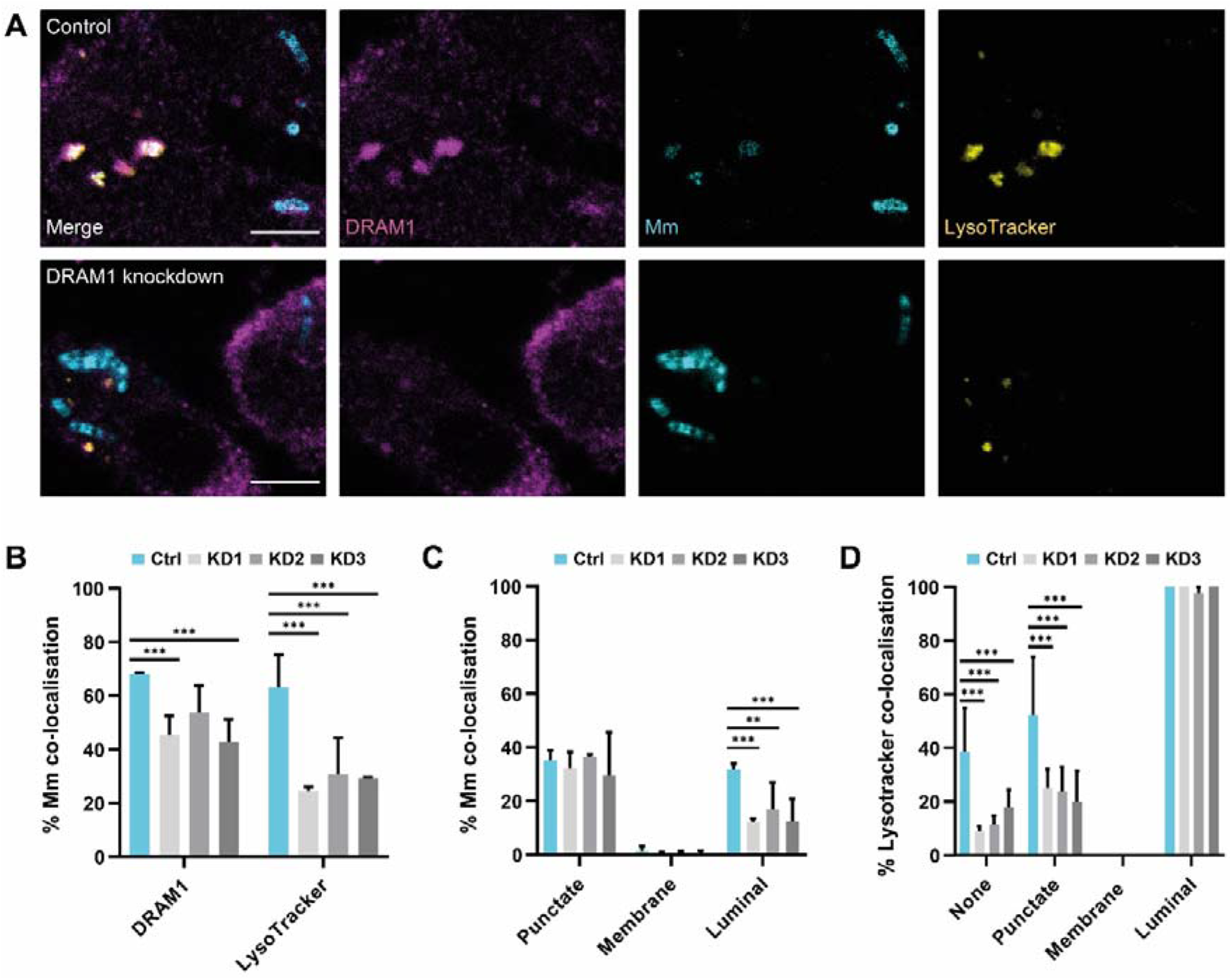
DRAM1 knockdown reduces acidification of Mm-containing vesicles. (A) Representative examples of DRAM1 (magenta) and LysoTracker (yellow) colocalisation with Mm (cyan) n control (Ctrl) and DRAM1 knockdown cell lines (KD1-3) at 120 min post infection Scale bars: 5 μm (B) Percentage of colocalisation of Mm with DRAM1 or LysoTracker in DRAM1 knockdown and control cell lines. (C) Percentage of colocalisation of Mm with DRAM1 in punctate, membrane, and luminal patterns in DRAM1 knockdown and control cell lines. (D) Percentage of colocalisation of LysoTracker with DRAM1-negative Mm (none) and DRAM1-positive punctate, membrane and luminal Mm patterns in DRAM1 knockdown and control cell lines. Data are accumulated from two independent experiments (nCtrl=117, nKD1=107, nKD2=92, nKD3=93 bacterial clusters). Statistical significance is assessed by logistic regression and pairwise comparison with Turkey correction. (*p<0.05; **p<0.01; ***p<0.001; ****p<0.0001)

### 3.4 DRAM1 mediates LC3 trafficking to Mm

Because we identified that LC3 is recruited to Mm-containing vesicles (Fig.3), we next wanted to assess if this step in the autophagy process is mediated by DRAM1. We found that the overall percentage of LC3-Mm colocalisation was reduced in all knockdown cell lines, similar to the overall percentage of LysoTracker-Mm colocalisation (Fig. 5A,B). Compared to the control, there was no significant reduction in punctate LC3-Mm colocalisation (Fig.5C), though the punctate LC3-Mm clusters did display reduced colocalisation with LysoTracker (Fig.5D). DRAM1 deficiency did not reduce basal LC3 levels, as shown by immunostaining and western blot analysis (Supplementary Fig.2 D,E,F,J); therefore this could not explain the decreased percentage of colocalisation between Mm and LC3. While the punctate LC3-Mm colocalisation was not affected, the DRAM1 knockdown cell lines showed a reduction of the luminal LC3-Mm colocalisation pattern (Fig.5C). The near-complete colocalisation of luminal LC3-Mm patterns with LysoTracker remained unchanged with DRAM1 knockdown (Fig.5D). To conclude, these results support that DRAM1 promotes the trafficking of LC3 to Mm and provide further evidence for the role of DRAM1 in acidification of Mm-containing vesicles.

**Figure 5.**
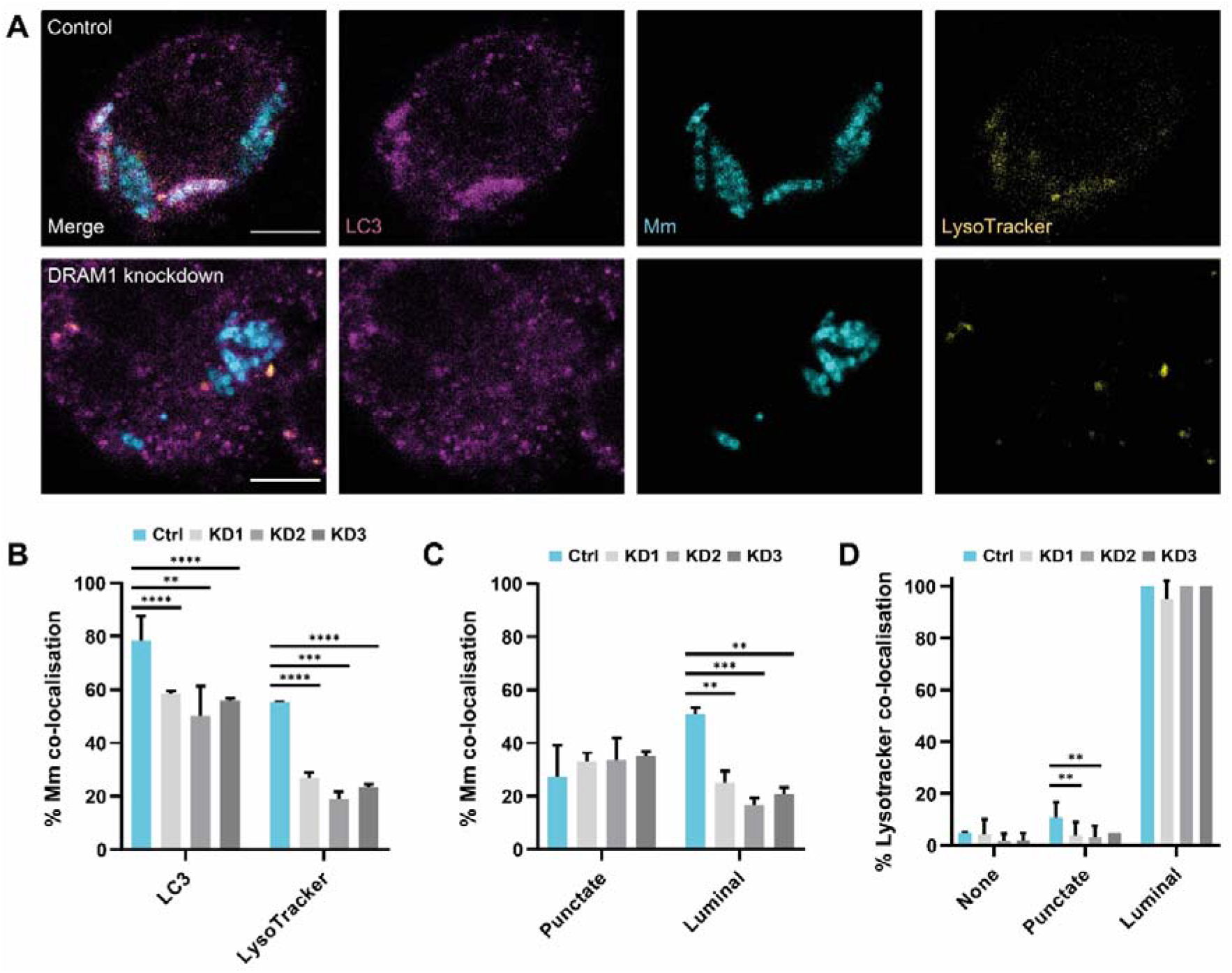
DRAM1 knockdown reduces LC3 trafficking to Mm. (A) Representative examples of LC3 (magenta) and LysoTracker (yellow) colocalisation with Mm (cyan) in DRAM1 knockdown (KD1-3) and control (Ctrl) cell lines at 120 min post infection. Scale bars: 5 μm (B) Percentage of colocalisation of Mm with LC3 or LysoTracker in DRAM1 knockdown and control cell lines. (C) Percentage of colocalisation of Mm with LC3 in punctate and luminal patterns in DRAM1 knockdown and control cell lines. (D) Percentage of colocalisation of LysoTracker with LC3-negative Mm and LC3-positive punctate and luminal Mm patterns in DRAM1 knockdown and control cell lines. Data are accumulated from two independent experiments (nCtrl=210, nKD1=184, nKD2=128, nKD3=120 bacterial clusters). Statistical significance is assessed by logistic regression and pairwise comparison with Turkey correction. (*p<0.05; **p<0.01; ***p<0.001; ****p<0.0001)

### 3.5 DRAM1 promotes the fusion of LAMP1-positive lysosomes with Mm-containing vesicles

The reduced acidification of Mm-containing vesicles under DRAM1 knockdown conditions prompted us to investigate the role of DRAM1 in mediating the interaction between Mm and the lysosomal marker protein LAMP1. Similar to the DRAM1 immunostaining definitions, we could classify LAMP1-Mm associations in three patterns: membrane, punctate and luminal (Fig.6A). Consistent with the reduced LysoTracker staining, we observed a reduction in the overall level of LAMP1-Mm colocalisation as a result of DRAM1 knockdown (Fig.6B,C). DRAM1 knockdown reduced the luminal LAMP1-Mm colocalisation patterns but did not reduce the membrane or punctate LAMP1-Mm patterns (Fig.6D). Furthermore, DRAM1 knockdown reduced the colocalisation of membrane LAMP1-Mm clusters with LysoTracker, but not the punctate or luminal patterns (Fig.6E). Based on immunostaining and western blot analysis of non-infected cells, the effect of DRAM1 knockdown on LAMP1-Mm patterns is not caused by a general reduction in LAMP1 protein levels (Supplementary Fig.3 C,D,G). In conclusion, the prominent effect of DRAM1 knockdown on the luminal LAMP1-Mm patterns is consistent with a role for DRAM1 in mediating the fusion of lysosomes with Mm-containing vesicles.

**Figure 6.**
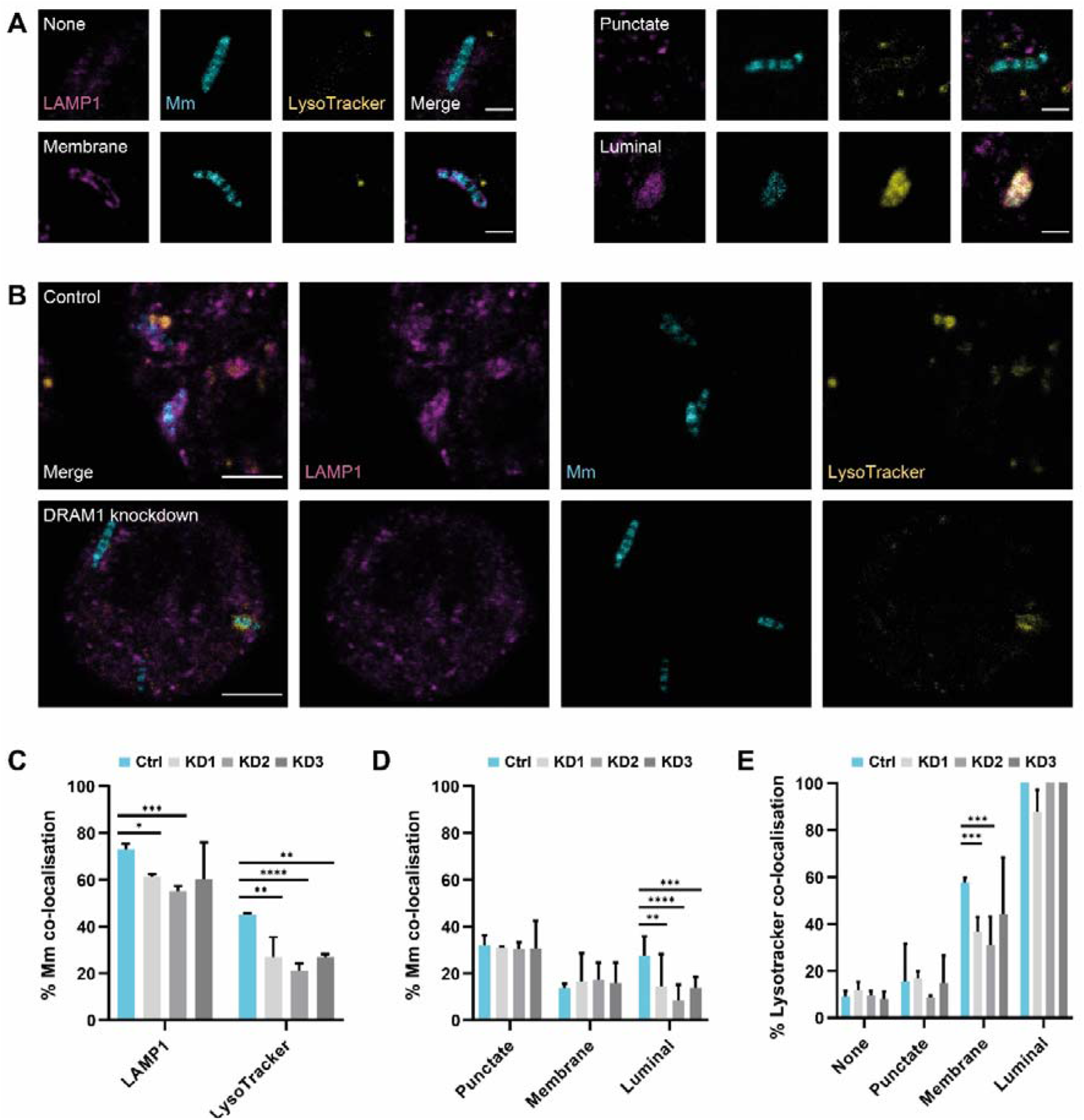
DRAM1 knockdown reduces fusion between lysosomes and Mm-containing vesicles. (A) Representative examples of LAMP1 (magenta) colocalisation patterns with Mm (cyan), referred to as membrane, punctate, and luminal patterns, at 120 min post infection. Acidification of Mm-containing vesicles is assessed by LysoTracker staining (yellow). Scale bars: 2 μm (B) Representative examples of LAMP1 (magenta) colocalisation with Mm (cyan), and with acidification of Mm-containing vesicles assessed by LysoTracker staining (yellow) in DRAM1 knockdown (KD1-3) and control cell lines at 120 min post infection. Scale bars: 5 μm (C) Percentage of colocalisation of Mm with LAMP1 or LysoTracker in DRAM1 knockdown and control cell lines. (D) Percentage of colocalisation of Mm with LAMP1 in punctate, membrane, and luminal patterns in DRAM1 knockdown and control cell lines. (E) Percentage of colocalisation of LysoTracker with LAMP1-negative Mm and LAMP1-positive Mm punctate, membrane, and luminal Mm patterns in DRAM1 knockdown and control cell lines. Data are accumulated from two independent experiments (nCtrl=156, nKD1=126, nKD2=153, nKD3=118 bacterial clusters). Statistical significance is assessed by logistic regression and pairwise comparison with Turkey correction. (*p<0.05; **p<0.01; ***p<0.001; ****p<0.0001)

### 3.6 DRAM1 is required for macrophage defence against Mm

Having determined the requirement of DRAM1 for Mm colocalisation with LysoTracker, LC3 and LAMP1, we wished to understand whether this role of DRAM1 in Mm vesicle trafficking impacts the susceptibility of RAW 264.7 macrophages to Mm infection. DRAM1 knockdown cell lines were infected with Mm for 1 hour to understand whether DRAM1 affected phagocytosis. The results showed no significant difference in the percentage of infected cells between the knockdown and control groups, indicating that DRAM1 does not impair the ability of RAW264.7 macrophages to phagocytose Mm (Fig7.A,B). Next, to determine if DRAM1 reduces Mm burden, cells were cultured for a total of 7 hours. Results showed that the percentage of infected cells in DRAM1 knockdown cell lines was higher than in the control group at 7 hours post infection (Fig7.C,D), indicating that DRAM1 protects against Mm infection without affecting phagocytosis in RAW 264.7 macrophages.

**Figure 7.**
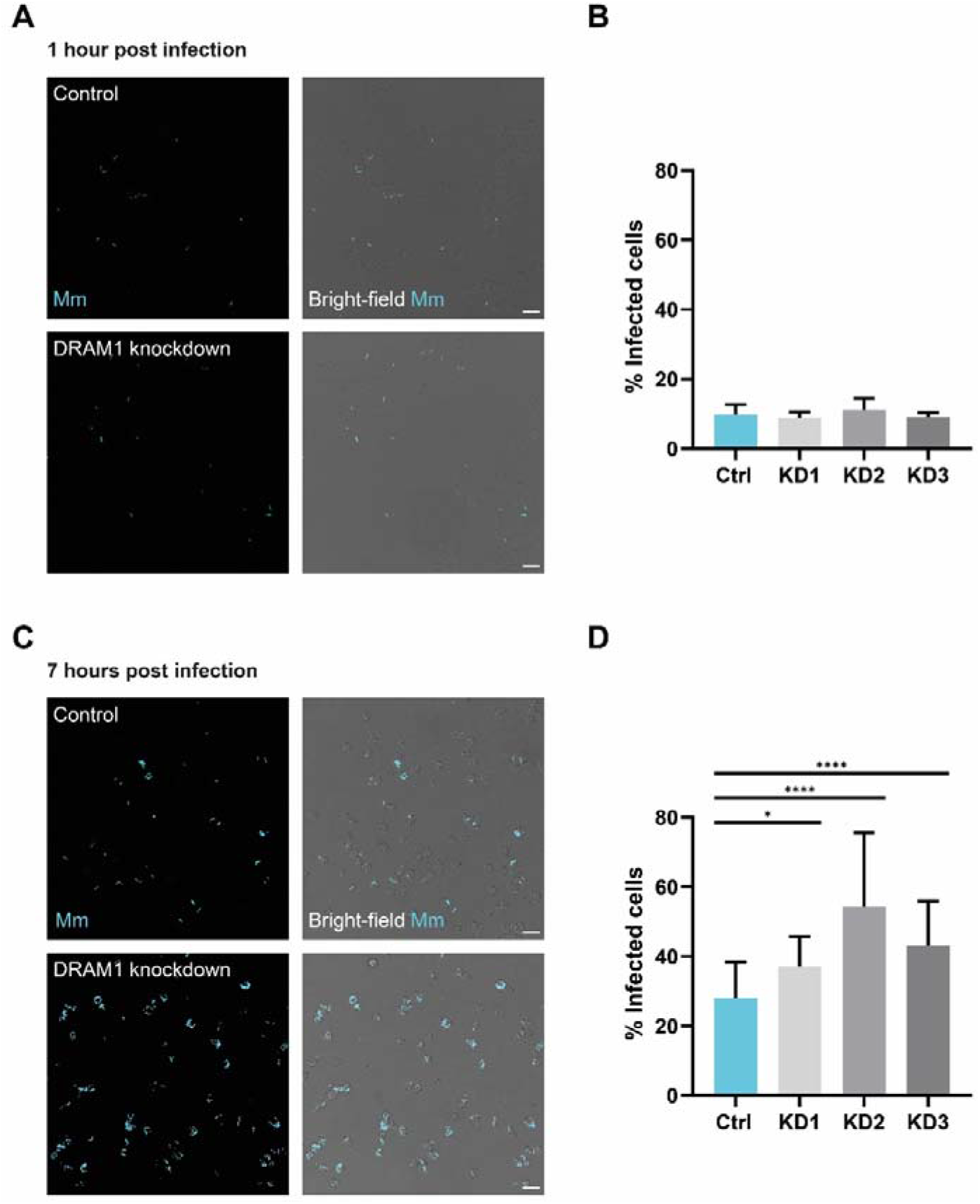
DRAM1 knockdown increases the susceptibility of macrophages to Mm infection. (A,C) Representative examples of infected cells in DRAM1 knockdown and control cells at 1 h (A) and 7 h (C) post infection. Scale bars: 20 μm (B, D) Percentage of infected cells in DRAM1 knockdown (KD1-3) and control cells at 1 h (B) and 7 h (D) post infection. Data are accumulated from three independent experiments with 12 images each. Statistical significance is assessed by logistic regression and pairwise comparison with Turkey correction. (*p<0.05; **p<0.01; ***p<0.001; ****p<0.0001)

## 4. Discussion

Understanding the mechanisms that deliver pathogens to lysosomes for degradation is crucial for developing novel therapeutic interventions against pathogens such as mycobacteria that are able to counteract both the phagolysosomal pathway and autophagy [37]. Based on studies in the zebrafish model, DRAM1 has emerged as a host resistance factor that promotes the delivery of mycobacteria to autophagosomes and lysosomes [18,27]. Here we confirm and extend these findings in a mammalian model, demonstrating the colocalisation that occurs between DRAM1 and mycobacteria in the context of autophagosomal and lysosomal compartments during the course of Mm infection in RAW264.7 macrophages. Furthermore, we show that DRAM1 knockdown impairs the macrophage defence response against Mm infection. These results support the central role of DRAM1 in the lysosomal delivery of mycobacteria.

We have previously detected DRAM1 colocalisation with Mtb in primary human macrophages but this cell type did not permit efficient DRAM1 knockdown studies [18]. Knockdown as well as loss-of-function mutation analysis of zebrafish Dram1 supported its requirement for autophagic defence against Mm [18,27]. However, a detailed analysis of DRAM1-mycobacteria colocalisation as well as a functional study in a mammalian cell type were still lacking. Therefore, we now employed an Mm infection model in murine RAW 264.7 macrophages. In uninfected RAW 264.7 cells, we detected DRAM1 on lysosomes, which is consistent with studies in other cell types that identified DRAM1 and its splice variants as largely endosomal/lysosomal proteins although being present also on autophagosomes, endoplasmic reticulum and peroxisomes [21,36]. In addition, we found that DRAM1 was prominently expressed in regions at or near the plasma membrane of RAW264.7 macrophages. Plasma membrane expression of DRAM1 has also been observed in hepatocytes and two other members of the DRAM family, DRAM3 and DRAM5, also display plasma membrane expression [24,26,38]. Additionally, we observed the formation of circular DRAM1-positive structures near the plasma membrane that might be the result of endocytosis, for example of cell debris, and therefore derived from the plasma membrane in regions where DRAM1 is highly concentrated. This expression pattern of DRAM1 is particularly interesting in the light of a recent study reporting a role for DRAM1 in extracellular vesicles formation [38]. Since exosomes are crucial for communication between immune cells, this might point to an unexplored role for DRAM1 in the intercellular propagating of signals during infection.

In the present work we focused on the role of DRAM1 in intracellular host defence. During Mm infection, we found that DRAM1 signal occasionally enveloped mycobacteria in a circular pattern, but most frequently appeared in a punctate pattern closely associated with mycobacteria, which gradually progressed to a full colocalisation pattern. The progression from partial to full colocalisation was concomitant with DRAM1-LC3-mycobacteria colocalisation as well as DRAM1-LysoTracker-mycobacteria colocalization and DRAM1-LAMP1-mycobacteria colocalisation, which strengthens the previous functional studies in zebrafish that linked Dram1 to the trafficking of mycobacteria along the (auto)phagolysosomal pathway [18,27]. Based on the colocalisation patterns and their frequency over time, we propose the following sequence of events. After phagocytosis, Mm first resides in a DRAM1-negative non-acidic phagosome and subsequently DRAM1 may be recruited to the phagosomal membrane. Lysosomes containing DRAM1 within the membrane may then fuse with the Mm-containing vesicle, leading to its acidification. Alternatively, Mm escapes from the phagosome into the cytoplasm and is subsequently trapped in an autophagosome. Continued recruitment of DRAM1 to Mm-containing vesicles of either phagosomal or autophagosomal nature would result in an increase in partial DRAM1-mycobacteria colocalisation. Subsequently, we hypothesise that DRAM1 becomes luminal following fusion between phagosomes or autophagosomes with lysosomes, resulting in full DRAM1-mycobacteria colocalisation in the acidic and degradative environment of autolysosomes or endolysosomes. Another potential explanation for the full colocalisation pattern is that it is derived from the invagination of the membrane enclosing Mm, resulting in the formation of multi-vesicular bodies (MVBs) that contain intraluminal vesicles with DRAM1 in the membrane. This would imply that lysosomes do not fuse until after complete DRAM1-mycobacteria colocalisation, as MVBs are formed at a late stage of vesicle maturation before lysosome fusion [39].

Previously, DRAM1 was shown to regulate autophagic flux through the lysosomal v-ATPase in human A549 cells under conditions of mitochondrial stress [40]. To determine DRAM1’s function in the (auto)phagolysosomal pathway of infected macrophages, we generated shRNA knockdown RAW 264.7 cell lines. DRAM1 knockdown led to reduced LC3-association, as well as reduced acidification of Mm-containing vesicles, suggesting that DRAM1 mediates autophagic targeting and acidic vesicle trafficking to Mm. This reflects results seen in zebrafish, where dram1 knockdown or mutation resulted in reduced GFP-LC3-Mm colocalisation and reduced LysoTracker-Mm colocalisation [18,27]. Extending from this, our analysis in RAW 264.7 macrophages demonstrated that there was reduced acidification of Mm-containing compartments showing partial or no colocalisation with the remaining DRAM1 protein in the knockdown lines. In contrast, despite DRAM1 knockdown, Mm-containing vesicles that showed full colocalisation with DRAM1 remained acidic. This suggests that DRAM1 affects trafficking of acidic vesicle to Mm-containing vesicles primarily at the early stages of vesicle maturation. In agreement, the effect of DRAM1 knockdown on LysoTracker-Mm colocalisation was larger than the effect on LAMP1-Mm colocalisation, suggesting that DRAM1 knockdown affects not only the fusion of lysosomes with Mm-containing vesicles but also the acidification of these vesicles due to fusion with late endosomes as an earlier step in the maturation process. The proposed role of DRAM1 in promoting fusion between acidic vesicles and Mm-containing compartments is supported by results observed in zebrafish overexpressing Dram1, where large vesicles containing membrane remnants were observed in transmission electron micrographs, suggesting that DRAM1 mediates multiple vesicle fusion events [18]. The stimulatory action of DRAM1 on vesicle trafficking and fusion in the (auto)phagolysosomal pathway likely enhances the microbicidal activity against mycobacteria, as we observed that DRAM1 knockdown led to an overall increased Mm infection burden in RAW 264.7 macrophages.

## 5. Conclusions

The results of this study, which show the requirement of DRAM1 for antibacterial autophagy and lysosomal delivery of Mm in RAW 264.7 macrophages, support the role of DRAM1 as a host resistance factor that is conserved across zebrafish and mammals. Therefore, DRAM1 represents a putative target for host-directed anti-mycobacterial therapy that could help overcome current challenges posed by anti-microbial resistance. Targeting DRAM1 may find wider applicability against intracellular pathogens, considering our recent results that DRAM1 also protects against infections with Salmonella enterica serovar Typhimurium and Aspergillus fumigatus [41,42]. Further elucidating its mechanism of action will facilitate the development of therapeutic strategies that exploit the role of DRAM1 in vesicle trafficking and fusion in the (auto)phagolysosomal pathway.

## Acknowledgements

The authors thank Gerda Lamers and Joost Willemse for help with microscopy analysis, Patrick van Hage for advice on statistical analysis, Sander van Kasteren (Leiden Institute of Chemistry) for the RAW264.7 cell line, Kevin Takaki (Department of Microbiology, Uni-versity of Washington, USA) for the Mm M-strain and Martijn Rabelink (Leiden University Medical Center) for shRNA constructs. JX was supported by a fellowship from the China Scholarship Council, MV by grant VI.Veni.192.151 from the Dutch Research Council (Nederlandse Organisatie voor Wetenschappelijk Onderzoek, NWO), and M.v.d.V. and A.H.M. by NWO-TTW grant 13259 from the NWO Domain Applied and Engineering Sciences.

## Author contributions

Conceptualization, A.H.M. and M.v.d.V.; methodology, M.V., A.H.M and M.v.d.V.; formal analysis, A.B.K. and J.X.; investigation, A.B.K., J.X., S.A.G.E. and C.S.; writing—original draft preparation, A.B.K. and J.X.; writing—review and editing, A.H.M and M.v.d.V.; visualization, A.B.K. and J.X.; supervision, M.V., A.H.M and M.v.d.V.; funding acquisition, J.X., M.V. and A.H.M. All authors have read and agreed to the published version of the manuscript.

## Supplementary figures

**Supplementary figure 1.**
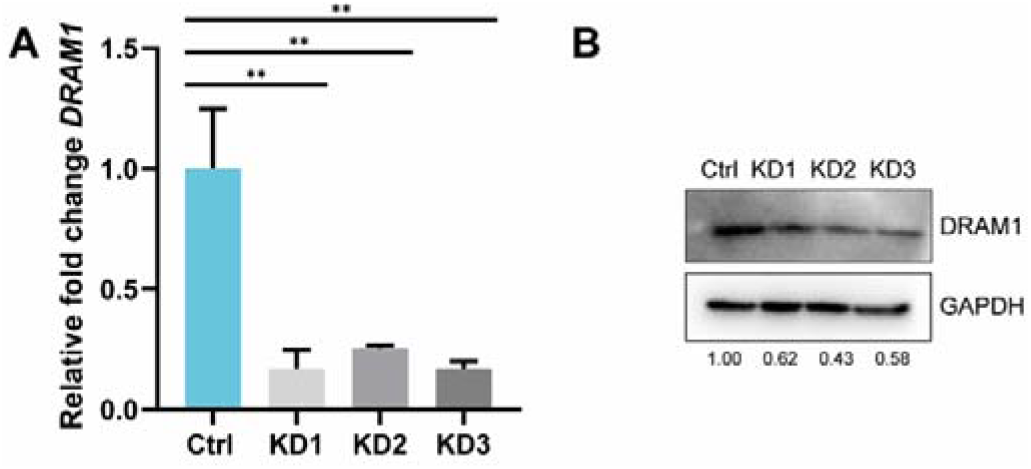
Confirmation of DRAM1 knockdown in RAW 264.7 macrophages. (A) qPCR analysis of Dram1 expression levels in control and knockdown cell lines (KD1-3). Statistical significance is assessed by logistic regression and pairwise comparison with Turkey correction. (**p<0.01) (B). Western blot analysis of DRAM1 protein levels in control and knockdown cells. Each cell line is analyzed in duplicate. DRAM1 protein levels relative to the control and normalized to GAPDH are indicated.

**Supplementary figure 2.**
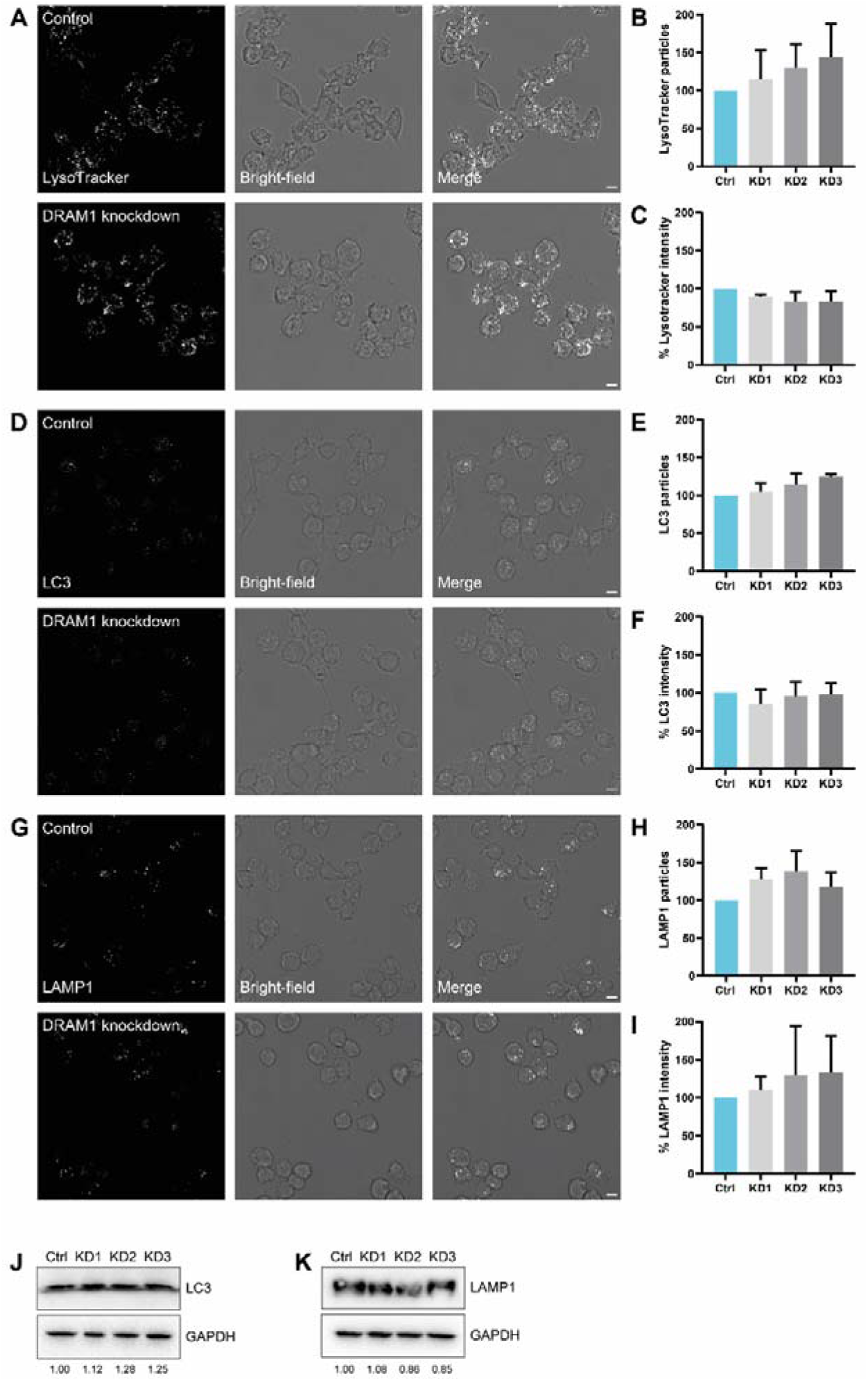
DRAM1 knockdown does not affect overall levels of LysoTracker, LC3 and LAMP1. Data are accumulated from three independent experiments with 12 images each and particles/intensity analysis are normalized by number of cells. (A,D,G) Representative examples of LysoTracker (A), LC3 (D) and LAMP1 (G) staining in non-infected DRAM1 knockdown (KD1-3) and control cells. Scale bars: 5 μm (B,C,E,F,H,I) Relative quantification of LysoTracker (B,C), LC3 (E,F) and LAMP1 (H,I) particles and signal intensity in noninfected DRAM1 knockdown and control cells.(J,K) Western blot analysis of LC3 (J) and LAMP1 (K) protein level relative to GAPDH in noninfected DRAM1 knockdown and control cells.

